# Pulsation changes link to impaired glymphatic function in a mouse model of vascular cognitive impairment

**DOI:** 10.1101/2021.06.08.447375

**Authors:** Mosi Li, Akihiro Kitamura, Joshua Beverley, Juraj Koudelka, Jessica Duncombe, Bettina Platt, Ulrich K. Wiegand, Roxana O. Carare, Rajesh N. Kalaria, Jeffrey J. Iliff, Karen Horsburgh

## Abstract

Large vessel disease and carotid stenosis are key mechanisms contributing to vascular cognitive impairment (VCI) and dementia. Our previous work, and that of others, using rodent models, demonstrated that bilateral common carotid stenosis (BCAS) leads to cognitive impairment via gradual deterioration of the glial-vascular unit and accumulation of amyloid-β (Aβ) protein. Since brain-wide drainage pathways (glymphatic) for waste clearance, including Aβ removal, have been implicated in the pathophysiology of VCI via glial mechanisms, we hypothesized that glymphatic function would be impaired in a BCAS model and exacerbated in the presence of Aβ. Male wild-type and Tg-SwDI (model of microvascular amyloid) mice were subjected to BCAS or sham surgery which led to a reduction in cerebral perfusion and impaired spatial learning and memory. After 3 months survival, glymphatic function was evaluated by cerebrospinal fluid (CSF) fluorescent tracer influx. We demonstrated that BCAS caused a marked regional reduction of CSF tracer influx in the dorsolateral cortex and CA1-DG molecular layer. In parallel to these changes increased reactive astrogliosis was observed post-BCAS. To further investigate the mechanisms that may lead to these changes, we measured the pulsation of cortical vessels. BCAS impaired vascular pulsation in pial arteries in WT and Tg-SwDI mice. Since our findings show that BCAS may influence VCI by impaired glymphatic drainage and reduced vascular pulsation we propose that these additional targets need to be considered when treating VCI.

## Introduction

Cerebral vascular disease (CVD) is a major contributor to vascular cognitive impairment (VCI) and dementia such as Alzheimer’s disease (Gorelick *et al.*, 2011; Montine *et al.*, 2014). Large well-characterized cohort studies have highlighted the co-existence of vascular disease with Alzheimer’s disease (De Jong *et al.*, 1997; de la Torre, 2000a,b,c; Esiri *et al.*, 1999; Hachinski and Munoz, 1997; Snowdon *et al.*, 1997). Key neuroimaging features (white matter lesions, microbleeds, lacunes and perivascular spaces) are found in both Alzheimer’s disease and VCI sharing a number of vascular risk factors, such as hypertension, diabetes and atherosclerosis (Dichgans and Leys, 2017). Vascular risk factors in midlife are also associated with increased burden of Alzheimer-related pathology, such as amyloid protein, suggesting a direct relationship (Gottesman *et al.*, 2017).

Chronic cerebral hypoperfusion has been proposed as a central common mechanism which contributes to cognitive decline and degenerative processes leading to dementia (Duncombe *et al.*, 2017). Global reductions in blood flow are associated with increased risk of progression from mild cognitive impairment to dementia suggesting that perfusion plays a key role in disease progression (Arbel-Ornath *et al.*, 2013; Chao *et al.*, 2010). Reduced cerebral perfusion has been linked to white matter attenuation, a key feature common to both Alzheimer’s disease and dementia associated with CVD (Barker *et al.*, 2014, Schuff *et al.*, 2009). Common artery stenosis of varying degrees is invariably associated with cognitive impairment (Alosco *et al.*, 2013; Balestrini *et al.*, 2013; Cheng *et al.*,2014; Cheng *et al.*, 2012; Johnston *et al.*, 2004) and carotid stenosis (>25%) has been linked to a greater burden of white matter hyperintensities (Romero et al,, 2009). Large and small vessel disease is also linked to Alzheimer’s disease dementia (Arvanitakis *et al.*, 2016). Reduced cerebral perfusion, impaired cerebrovascular reactivity and hemodynamic responses are increasingly recognized in the early stages of Alzheimer’s disease (de la Torre, 2012b; Hughes *et al.*, 2014). Our work and others using animal models have shown that chronic cerebral hypoperfusion as a result of bilateral carotid stenosis leads to cognitive decline through mechanisms that involve hypoxia-induced white matter damage and gradual deterioration of the neuro-glial-vascular unit including endothelial dysfunction, microvascular inflammation and BBB leakage (Fowler *et al.*, 2017; Holland *et al.*, 2011; Kitamura *et al.*, 2017; Roberts *et al.*, 2018; Shibata *et al.*, 2004). There is substantial evidence that reduced blood flow contributes to vascular disease. However, a causal relationship remains a matter of controversy largely due to the cross-sectional nature of clinical studies and, in the few longitudinal studies conducted, reduced blood flow occurs subsequent to vascular disease burden (de la Torre, 2012a). Impaired glymphatic function is emerging as a key player in vascular disease and dementia. The glymphatic pathway is a brain wide clearance process that relies on the movement of CSF along the perivascular network facilitated by aquaporin-4 water channels on the astroglial endfeet to promote the elimination of waste out of the brain (Iliff *et al.*, 2012). CSF flow within the perivascular space (PVS) is regulated by cerebrovascular pulsatility and constriction, which is now considered to be a key factor regulating glymphatic function (Iliff *et al.*, 2013; Mestre *et al.*, 2018, 2020). Enlarged PVS, identified by neuroimaging, are a common feature of cerebral vascular disease and dementia linked to vascular risk factors and inflammation (Aribisala *et al.*, 2014; Ding *et al.*, 2017; Doubal *et al.*, 2010; Potter *et al.*, 2015; Shi and Wardlaw, 2016; Wardlaw *et al.*, 2013). There is also evidence of impaired glymphatic function in pre-clinical models relevant to CVD. Notably advanced age, acute ischemic stroke and multi-infarct stroke, diabetes and subarachnoid haemorrhage (SAH) have all been shown to have a major impact on glymphatic drainage (Gaberel *et al.*, 20104; Kress *et al.*, 2014; Wang *et al.*, 2017). Disturbances of the glymphatic function are also related to a build-up of Aβ in both human and rodent brain (Shokri-Kojori *et al.*, 2018; Xu *et al.*, 2015).

Recent studies from our group and others have shown that carotid stenosis reduces cerebral perfusion and alters the Aβ peptide pools culminating in cerebral amyloid angiopathy (CAA) and vascular related lesions (Okamoto *et al.*, 2012; Salvadores *et al.*, 2017). In light of the evidence that flow-limiting large-vessel stenosis contributes to vascular and Alzheimer’s disease pathophysiology (Gupta and Iadecola, 2015), and that impaired glymphatic function is a key contributor to impaired Aβ clearance we hypothesized that the complex interaction of Alzheimer’s disease and carotid stenosis leading to cognitive impairment occurs via impaired glymphatic function in addition to perfusion deficits. We interrogated this by examining glymphatic influx in a well-characterized murine model of vascular cognitive impairment induced by bilateral common carotid stenosis (BCAS) (Shibata *et al.*, 2004) and then assessed Aβ accumulation in a model of microvascular amyloid (Tg-SwDI) post-BCAS. We further assessed astrocytes and cerebral vascular pulsation as potential mechanisms since they govern CSF-ISF exchange in murine brain (Iliff *et al.*, 2013).

## Materials and Methods

### Mice

All experiments were conducted in accordance with the UK Home Office Animals (Scientific Procedures) Act 1986 and additional local ethical and veterinary approval (Biomedical Research Resources, University of Edinburgh) and the ARRIVE guidelines. We used male C57Bl/6J (Charles River Laboratories Inc, UK) and Tg-SwDI mice (transgenic mice with Swedish, Dutch and Iowa mutations in human amyloid precursor protein (APP), with primarily microvascular amyloidosis) for all experiments. At the outset, mice from cohort 1 (n=42) (Tg-SwDI and wild-type littermates at 7-9 months old) were randomly assigned to experiments of MRI and behavioural tests, and tissues were collected for evaluation of microvascular amyloid level. A second cohort of mice (cohort 2) (n=33) (Tg-SwDI at 5-7 months and imported C57Bl/6J mice at 4-5 months) were used for the investigation of glymphatic influx and astrogliosis. Mice from cohort 3 were used for in vivo investigation of vessel pulsation. Investigators were blinded to surgery and genotype throughout the data collection and analysis. Final group size for analysis: cohort 1: n=8 WT sham, n=10 WT BCAS, n=6 Tg-SwDI sham, n=10 Tg-SwDI BCAS. Cohort 2, n=10 WT sham, n=8 WT BCAS, n=7 Tg-SwDI sham, n=8 Tg-SwDI BCAS. Cohort 3, n=7 WT sham, n=7 WT BCAS, n=6 Tg-SwDI sham, n=7 Tg-SwDI BCAS.

### Bilateral common carotid stenosis (BCAS) surgery

BCAS surgery was performed under isoflurane anaesthesia by applying microcoils (0.18mm internal diameter, Sawane Spring Co, Shizuoka, Japan) permanently to both common carotid arteries. Details of surgical methods have been described in previous studies (Coltman *et al.*, 2011; Holland *et al.*, 2011; Reimer *et al.*, 2011; Shibata *et al.*, 2004). A 30-minute interval was given between two microcoils application to minimize the acute CBF changes caused by the placement of microcoils. Sham-operated animals underwent the identical procedure except the application of microcoils to both arteries. In cohort 1, one Tg-SwDI mouse was culled during surgery due to severe bleeding, after three days of surgery, two WT and five Tg-SwDI were culled due to poor recovery. Therefore, these mice were excluded from the study.

### Cerebral blood flow measure by arterial spin labelling

A 7.0T Agilen (Varian) preclinical MRI system was used to collect T1-weighted and arterial spin labelling (ASL) data. Experimental animals were anesthetized under 5% isoflurane in oxygen for induction then placed in an MRI compatible holder (Rapid Biomedical, Wurzburg, Germany). Isoflurane was maintained at 1.5% in oxygen during scanning. Rectal temperature was monitored and regulated at around 37 °C by an airflow heating system. Respiratory rate was regulated at 70- 100 breaths per minute. The T1-weighted images were acquired at 1.7 mm posterior to Bregma in stereotactic coordinates of Mouse Brain Atlas (Paxinos and Franklin, 2001). Resting cerebral blood flow was measured using ASL at the level corresponding to T1-weighted scans with a Look-Locker FAIR single gradient echo (LLFAIRGE) sequence. Maps of cerebral blood flow were constructed from ASL data in Matlab using in-house scripts. Cerebral blood flow maps were analysed in ImageJ (v1.46, NIH, Bethesda, MD, USA) using unbiased regions of interest from T1- weighted images acquired with the ASL sequence.

### Assessment of cognitive function using Barnes maze

A Barnes maze was used to assess the differences in spatial learning and memory at 3 months after BCAS or sham surgery (Barnes maze schedule shown in Figure 2A). The maze consists of one white circular platform and 20 circular holes around the outside edge of the platform, with 91.5 cm diameter and 115 cm height (San Diego Instruments). The maze was brightly lit with lamps and overhead room lights (450 lux), and an aversive white noise stimulus is played at 85 dB. There is one dark escape chamber attached to one of the holes allocated to each experimental animal. Visual cues were placed on the curtains and walls around the maze. There was one white cylinder with 10.5 cm diameter for retaining animals at the beginning of each trial. All the tests were recorded by a video-based automatic tracking system ANY-maze v 4.99. All the tests were performed in the behaviour testing room where the room temperature can be controlled at constant 20 °C.

### Acclimation and habituation

Animals were brought into the behavioural testing room and placed in the holding cylinder to acclimate to the testing environment for 10 seconds for two days before habituation. One week prior to the training session, animals were habituated to the maze and escape cage. Each mouse was placed in the holding cylinder for 10 seconds then allowed 3 minutes free exploration under low stress conditions after removal of the cylinder, without aversive white noise stimulation. Then mice were guided to the escape cage and allowed inside for 2 minutes. All the animals were allocated one fixed number for the cage during the behaviour test. The maze and the escape cage were cleaned with ethanol to avoid any olfactory cues between each trial.

### Visuo-spatial learning and working memory test (acquisition training)

During the training session, mice were trained to find the escape chamber over 6 days with 2 trials per day (60-minute inter-trial interval). The platform consists of 20 escape holes and the location of escape chamber remained constant to each mouse but was shifted clockwise 90 degree between mice to avoid any olfactory cues. The mouse was placed in the holding cylinder for 10 seconds.

The aversive white noise (85 dB) was given once the test started and switched off once the mouse entered the escape chamber. If the mouse failed to enter the target hole, the experimenter guided the mouse to the escape cage. The aversive stimulus was stopped as soon as the mouse entered the chamber.

### 72 hours probe

A probe trial was performed 72 hours after the final acquisition training and each mouse was allowed 90 seconds to explore the maze with the escape cage removed and the rest of elements remained same. The 72 hours probe trials aimed to test the long-term memory of the mice after a period of training to locate the escape chamber.

### Reversal training

During the reversal training session, mice were trained to find the escape cage following same procedure as the acquisition training phase, but with the allocated escape cage shifted 180 degree to the opposite side of the stage. The mice were trained over 3 days with 2 trials per day (60-minute inter-trial interval) in reversal training to evaluate the spatial learning ability in increased difficulty of task.

### Reversal probe

The reversal probe trial was performed 72 hours after the final reversal training. Animals were given 90 seconds to explore the maze with the escape cage removed. All the elements in reversal probe remained same as reversal training test.

### Measurements

Trials were recorded by a camera above the maze and measured using tracking software Any-maze version 4.99. Spatial learning was assessed by the total time to enter the escape cage (escape latency) and the total distance travelled (pathlength) during this period.

### Assessment of glymphatic function by intracisternal injection of fluorescent tracers

Mice were initially anaesthetized with isoflurane (5% in oxygen), then positioned on a stereotaxic frame and anaesthetic maintained at approximately 1.5% (in oxygen). The respiration was regulated using a ventilator. The posterior atlanto-occipital membrane was surgically exposed and a 32GA needle attached to a Hamilton syringe was inserted into cisterna magna. Dextran, fluorescein and biotin labelled 3 kDa soluble lysine fixable (D7156, Invitrogen) and ovalbumin Alexa Fluor® 594 conjugate 45 kDa (O34783, Invitrogen) tracers were mixed at 1:1 ratio and infused at a concentration of 5 μg/μl, at a rate of 0.5 μl/min over 20 minutes (10 μl total volume) through a syringe pump (Harvard Apparatus). The needle was held in place for 10 minutes and then removed, and atlanto-occipital membrane was sealed to avoid any reflux of CSF.

### Tissue processing

At the end of the experiments, mice from cohort 1 and 2 were transcardially perfused with 30 ml heparinized saline then whole brains were fixed in 4% paraformaldehyde in PBS for 24 hours. For cohort 1 1, brain tissues were further transferred into 30% sucrose solution in PBS for 72 hours. Brains were placed in pre-cool isopentane -42 °C for 5 minutes then stored in -80 °C freezer and coronal sections (12 µm) were cut using a cryostat. For cohort 2, the brains were sectioned into coronal planes (100 µm) on a vibratome then stored in cryoprotective medium in a -20 °C freezer.

### Imaging of fluorescent tracer movement

Tracer movement from the subarachnoid space of the cisterna magna into the brain was imaged using a slide scanner (ZEISS Axio Scan.Z1). Multi-channel whole-slice images of each animal at hippocampal level (-1.82 mm to bregma, Mouse Brain Atlas) was generated at 20x magnification. This included separate DAPI, Alexa Fluor 488 and Alexa Fluor 594 channels. All images were scanned using constant exposure time for each individual channel by the slide scanner. For the quantification of tracer movement into the brain, scanned images were analysed in ImageJ software (v1.46, NIH, Bethesda, MD, USA) as described previously (Iliff *et al.*, 2012) Region of interest (ROI) was defined using DAPI channel to identify anatomical regions. Auto-thresholding (triangle method) was used to measure the % area of positive signal that is the glymphatic CSF influx.

### Immunohistochemistry

Immunostaining was carried out according to standard protocols. Frozen sections were removed from the freezer and allowed to air dry for 30 minutes. Slides were washed in phosphate-buffered saline (PBS) followed by a series of ethanol (70%, 90% and 100%) for dehydration then placed in xylene for 10 minutes. Sections were rehydrated through serial ethanol (100%, 90% and 70%) then rinsed in water. Antigen retrieval was performed using 10mM citric buffer (PH 6.0) at 100 °C under pressure for 10 minutes then covered with proteinase K working solution for 10 minutes at room temperature. Sections were rinsed in PBS and incubated in blocking buffer (10% normal serum, 0.5% BSA) for 1 hour at room temperature. Subsequently, sections were incubated in primary antibody solution (amyloid 6E10, 1:1000, Covance, SIG-39320, mouse monoclonal antibody; COL4, 1:400, Fitzgerald, 70R-CR013X, rabbit polyclonal antibody) overnight at 4 °C. Sections were then rinsed in PBS and incubated in secondary antibody (anti-rabbit Alexa Fluor 546, 1:500, Invitrogen A-11010; anti-mouse Alexa Fluor 488 1:500, Invitrogen A-11001) for 1 hour at room temperature.

Vibratome sections were rinsed in PBS and mounted on to superfrost plus slides (VWR international) followed by serial ethanol (70%, 90% and 100%) and then placed in xylene for 10 minutes. Sections were rehydrated through serial ethanol (100%, 90% and 70%) then rinsed in running water. Antigen retrieval was performed using 10mM citric buffer (PH 6.0) at 100 °C under pressure for 10 minutes. Then sections were incubated in primary antibody solution (GFAP, 1:1000, Life technologies, 13-0300, Rat monoclonal antibody) overnight at 4 °C. Sections were rinsed in PBS and incubated in non-fluorescent biotinylated secondary antibody (anti-rat, 1:100, Vector Laboratories, YO809) for 1 hour at room temperature followed by 1 hour incubation with Vector ABC Elite kit (Vector Laboratories). Finally, sections were visualized with DAB peroxidase substrate kit (Vector Laboratories).

### Analysis of immunohistochemistry

Immunostained 12 µm frozen sections were analysed using a laser scanning confocal microscope (ZEISS LSM 710, Germany). Cortical amyloid load and blood vessel density were determined by measuring the percentage of areas occupied by 6E10 and COL4 staining, respectively. Vascular amyloid load was determined by colocalization analysis for blood vessels and amyloid by calculating the Mander’s coefficient and data shown as % vascular amyloid. Images from the cortex in Tg-SwDI mice were selected, amyloid, blood vessels (COL4) and vascular amyloid images were quantified at the brain regions from the pial surface to approximate the depth of 250 µm. Immunostained 100 µm thick vibratome sections were analysed using a slide scanner (ZEISS Axio Scan.Z1). Astrogliosis were assessed by measuring the percentage of stained area occupied by GFAP staining, using auto thresholding (triangle method). All measurements were carried out using ImageJ (v1.46, NIH, Bethesda, MD, USA)

### Cranial window implantation

Animals were initially anaesthetised using 4-5% isoflurane, delivered through a face mask via a ventilator and kept on 1.5-2% through the surgery. Subcutaneous injection of Caprofen (5 mg/kg) was administered at the start of the surgery. Body temperature was monitored through the surgery. Skin was removed to expose the skull, dried, and secured using VetBond (3M, #1649). Using a high-speed micro drill, an area of 6x3 mm was drilled over until a thin layer of bone was left. A drop of artificial cerebrospinal fluid (ACSF) (ACSF; 125 mM NaCl, 10 mM glucose, 10 mM HEPES, 3.1 mM CaCl2, 1.3 mM MgCl2, pH 7.4), was applied to the skull and left for 10 minutes. Using angled forceps, the skull was lifted without disrupting the dura. Hemocollagene (Septodont, #0459) soaked in ACSF is applied on the exposed brain for 5 minutes. A sterile coverslip was placed on top of the exposed brain, and secured by a mixture of liquid glue and dental cement. Immediately afterwards, a custom made head plate (Protolabs) was applied to the cranial window prep and secured by additional glue/cement mixture. The cranial window was left to dry for 5 minutes and the animal was placed for recovery. Animals were rested for 4 weeks prior to imaging.

### In vivo vascular pulsation assessment

In separate cohorts of WT and Tg-SwDI mice (cohort 3) cerebral vascular pulsatility was evaluated in WT and Tg-SwDI mice after sham and BCAS surgery.

Cerebral vascular pulsation was assessed through the cortical vascular network with the use of multiphoton microscopy (LaVision Biotech TriMScope with Nikon CFI-Apo 25x NA1.1 lens and Leica SP8 DIVE with IRAPO L 25x NA1.0 lens) using a method based on those described previously within the literature (Iliff *et al.*, 2013; Kress *et al.*, 2014). Cortical vascular network was visualized via multiphoton microscopy through the injection of fluorescently conjugated dextran into the blood stream (Rhodamine B, 20 mg/ml, Sigma R9379). The pulsatility of individual vessels was determined by positioning linescans orthogonal to the vessel’s axis. Linescans were performed at in a repeated loop scans at a frequency of 1086.96 Hz and 8000Hz to equal duration of 3600 ms. To calculate dynamic vessel width changes over time, vessel width (µm) within each region was plotted against time (ms). Vessel wall pulsatility (µm*ms) was calculated from the resulting graphs as the absolute value of area under the diameter-time plot, integrated to the running average calculated across the entire 3600 ms sampling time. To calculate the total number of dynamic vessel pulses (pulsation frequency) across the sampling time, the total number of peaks were also calculated from the same diameter-time plots.

### Statistical analysis

Data were analysed using a two-way ANOVA with surgery and genotype as two between-subject factors followed by Bonferroni’s multiple comparison test to compare CBF levels, CSF glymphatic drainage and astrogliosis. Statistical comparison of spatial learning was carried out by repeated measures ANOVA with surgery and genotype as between subject factors followed by Bonferroni’s multiple comparison test, the probe trials were carried out by using two-way ANOVA for comparison between groups. One sample t-test was used to compare the performance of each group with the chance. Mann Whitney U test was used to compare the amyloid burden, blood vessel density and pulsation. Statistical analysis was performed using IBM SPSS Statistics 22.

### Data availability

All relevant data are available in the manuscript and in the Supplementary Materials. Requests for materials should be addressed to K.H

## Results

### Regional cerebral perfusion is reduced post-BCAS in wild-type and Tg-SwDI mice

At the outset of the studies, we determined whether carotid stenosis affected cerebral blood flow (CBF) in Tg-SwDI compared to WT mice. Arterial spin labelling (ASL) showed the extent of reductions in regional CBF in the dorsolateral cortex and hippocampal CA1-DG region (Figure 1A). In the dorsolateral cortex (DL CTX), there was a significant main effect of surgery (F (1, 26) =14.816, p<0.001) but no effect in Tg-SwDI mice (p>0.05) on resting CBF (Figure 1B). Post-hoc analysis indicated that CBF was significantly reduced in BCAS mice in both wild-type (p=0.013) and Tg-SwDI (p=0.010) groups. Furthermore, in the hippocampal CA1-DG region, there was a main effect of surgery (F (1, 26) =17.96, p<0.001) but not genotype (p>0.05) (Figure 1C). Post-hoc analysis showed significantly reduced CBF in both wild-type (p=0.022) and Tg-SwDI (p=0.002) BCAS groups. Although there was a sustained and prominent reduction in CBF, there were no genotype differences indicating that perfusion was reduced to a similar extent in WT and Tg-SwDI mice.

**Figure 1:**
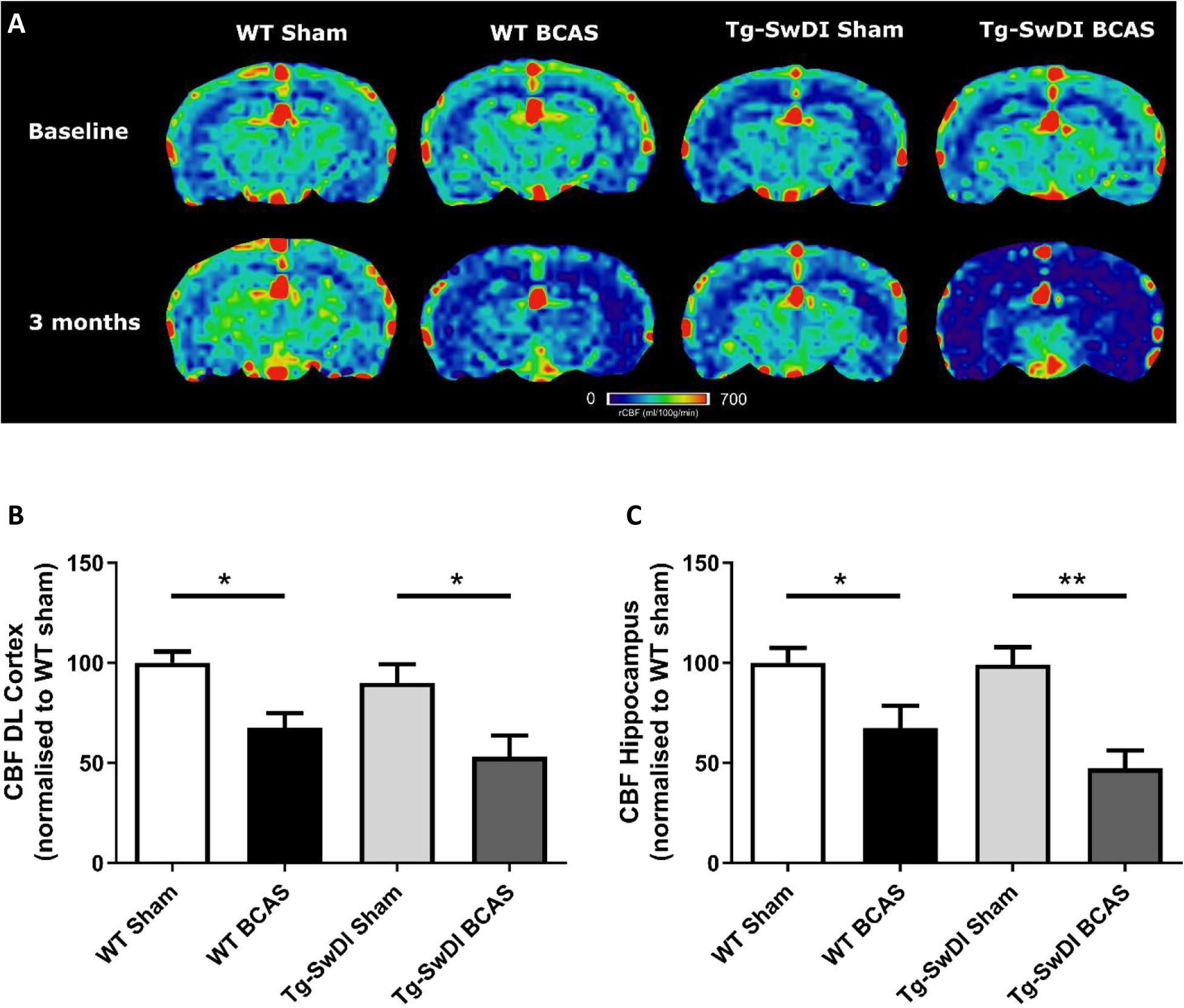
Decreased resting CBF following BCAS. MRI arterial spin labelling (ASL) was used to measure regional alterations in CBF. A Representative images of arterial spin labelling (ASL) from sham and BCAS WT and Tg-SwDI mice at 3 months following surgery. B and C A significant reduction of CBF in the brain cortex and hippocampus was determined post-BCAS but there was no genotype effect. Data are presented as mean ± SEM, n=6-10 per group

### BCAS causes a decline in spatial learning acquisition and cognitive flexibility in wild-type and Tg-SwDI mice

We and others have previously reported that BCAS caused short-term spatial working memory impairments and spatial learning and memory deficits in WT mice (Coltman *et al.*, 2011; Holland *et al.*, 2015; Kitamura *et al.*, 2017; Martin *et al.*, 2016; Patel *et al.*, 2017). In this study spatial learning and memory abilities were assessed in BCAS wild-type mice but additionally it was determined whether BCAS would cause an exacerbated impairment in Tg-SwDI mice. The Barnes maze paradigm was used to evaluate visuo-spatial learning whereby mice were trained to locate an escape hole using spatial cues over 6 days with 2 sessions per day (Figure 2A) and the escape latency measured. There was a significant effect of BCAS (F (1, 30) =9.60, p<0.01) but not genotype (p>0.05) on escape latency (Figure 2B). Given these measures might be affected by alterations in locomotion, we next analysed the distance travelled in the tests (pathlength) as additional measures to evaluate spatial learning. There was a significant effect of BCAS (F (1, 30) =5.826, p=0.022) but no effect of genotype (p>0.05) on the pathlength across groups (Figure 2B). To investigate the effect of BCAS on long-term memory, a probe test was taken after 72 hours of the final acquisition training to examine whether experimental animals remember the previous target location after removing the escape chamber. Data was quantified as the percentage of time each mouse spent in the target quadrant where the allocated chamber was previously located. WT sham (p=0.004), Tg-SwDI sham (p=0.021) and Tg-SwDI BCAS (p=0.025) all spent a significantly higher percentage of time than chance (25%) in the target quadrant but wild-type BCAS mice, did not perform above chance level (p=0.84) (Figure 2D). There was no significant effect of BCAS or genotype on the percentage of time spent in the correct quadrant across groups. Reversal training and probe trials were then undertaken to evaluate the ability of experimental animals to learn a new location and to test cognitive flexibility. The escape hole location was switched 180° to the opposite side of maze. The reversal probe was performed following 72 hours after the final reversal training trial. Only WT sham mice (p<0.05) spent a significantly higher percentage of time than chance in the target quadrant whereas all other groups did not perform above chance (WT BCAS, Tg-SwDI sham, Tg-SwDI BCAS p>0.05, respectively) (Figure 2E). Collectively the data demonstrate that BCAS impairs learning acquisition and cognitive flexibility.

**Figure 2:**
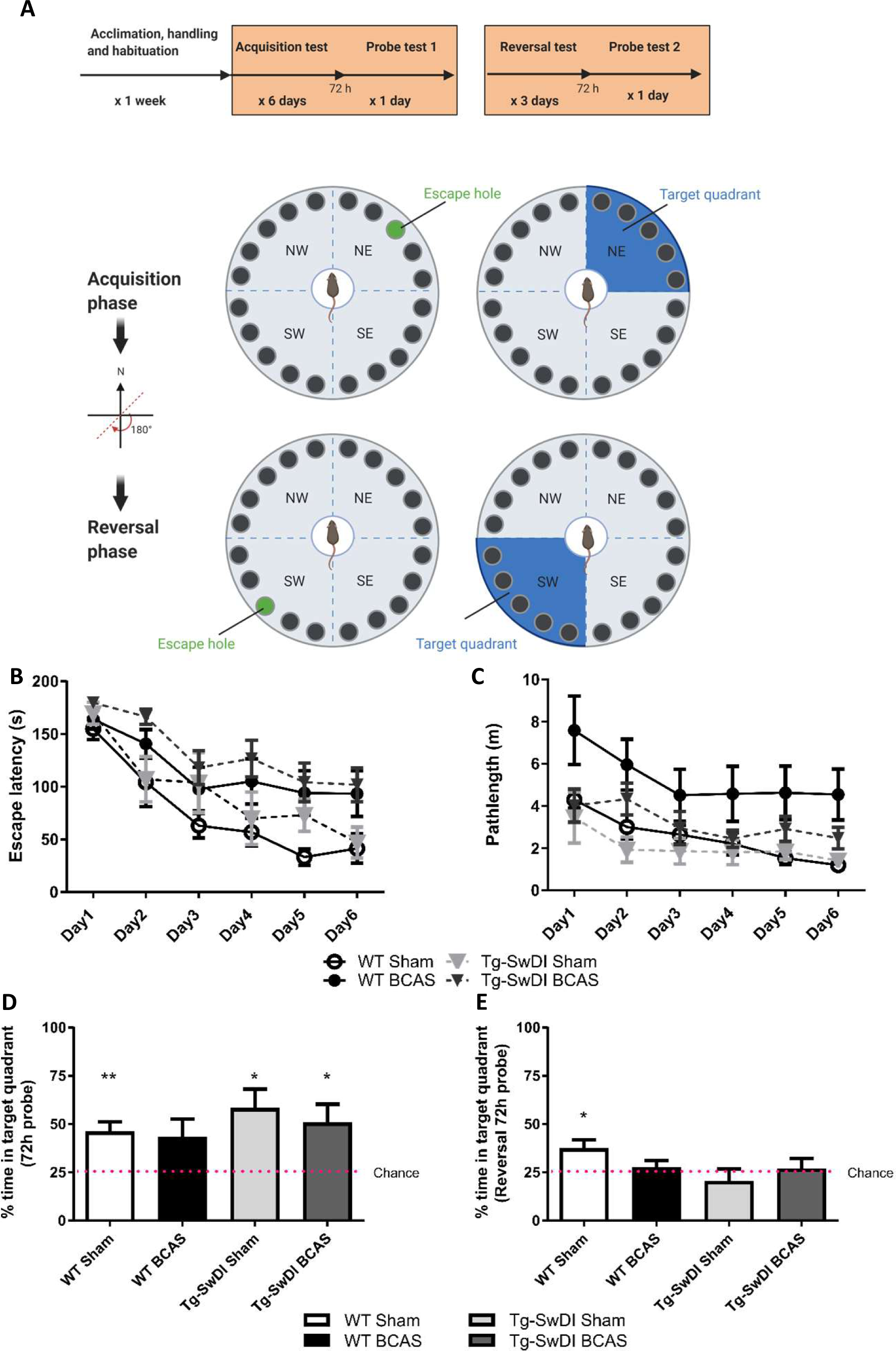
BCAS causes a decline in spatial learning acquisition and cognitive flexibility. Spatial learning and memory and cognitive flexibility were assessed using a Barnes maze at 3 months post-BCAS in WT and Tg-SwDI mice. A In the acquisition phase one hole (indicated by blue hole) was designated as the target hole with an escape box. A probe test was performed 72 hours after the last acquisition training session, in which the escape box was removed. In the reversal phase, the target hole was moved 180 degrees to the original target hole. Position opposite the original 1 day after the probe test. A reversal probe test was performed 72 hours after the last training session. B Spatial learning was assessed by comparing escape latency over 6 days with 2 sessions per day. There was a significant effect of BCAS surgery but not genotype across groups. C Pathlength measure was also used to evaluate spatial learning function. There was a significant effect of BCAS surgery but not genotype across groups. D In the acquisition 72h probe test all mice performed above chance except WT BCAS mice. E In the reversal probe test only WT sham mice performed above chance. Data are mean ± SEM, n=6-10 per group

### CSF glymphatic influx in the brain cortex

We next determined whether glymphatic function would be impaired post-stenosis at a time when both cerebral perfusion and cognitive abilities are impaired. To address this, we first examined the distribution of Evans blue following cisterna magna injection as a method to visualize glymphatic entry/influx and found that the dye distributed along the surface brain vessels (e.g. middle cerebral artery), along the superior sagittal sinus, inferior cerebral vein, and transverse sinus (Figure 3A). Following this CSF influx was then investigated by injection of fluorescently labelled CSF tracer (Dextran 3kDa, D-3) into the cisterna magna and tracer distribution evaluated by imaging ex vivo fixed brain slices co-labelled with markers of the basement membrane (COL4) and astrocytic end- feet (AQP4). CSF tracer influx was observed colocalized with the basement membrane and in the adjacent space (Figure 3B), colocalized with basement membrane (Figure 3C) and surrounded by astrocytic end-feet AQP4 (Figure S3). Intensity profile graphs show strong colocalization between CSF tracer (D-3) and vascular basement membrane (COL4) (Figure 3D and E) with partial tracer occupancy in the perivascular compartment (Figure 3D).

**Figure 3:**
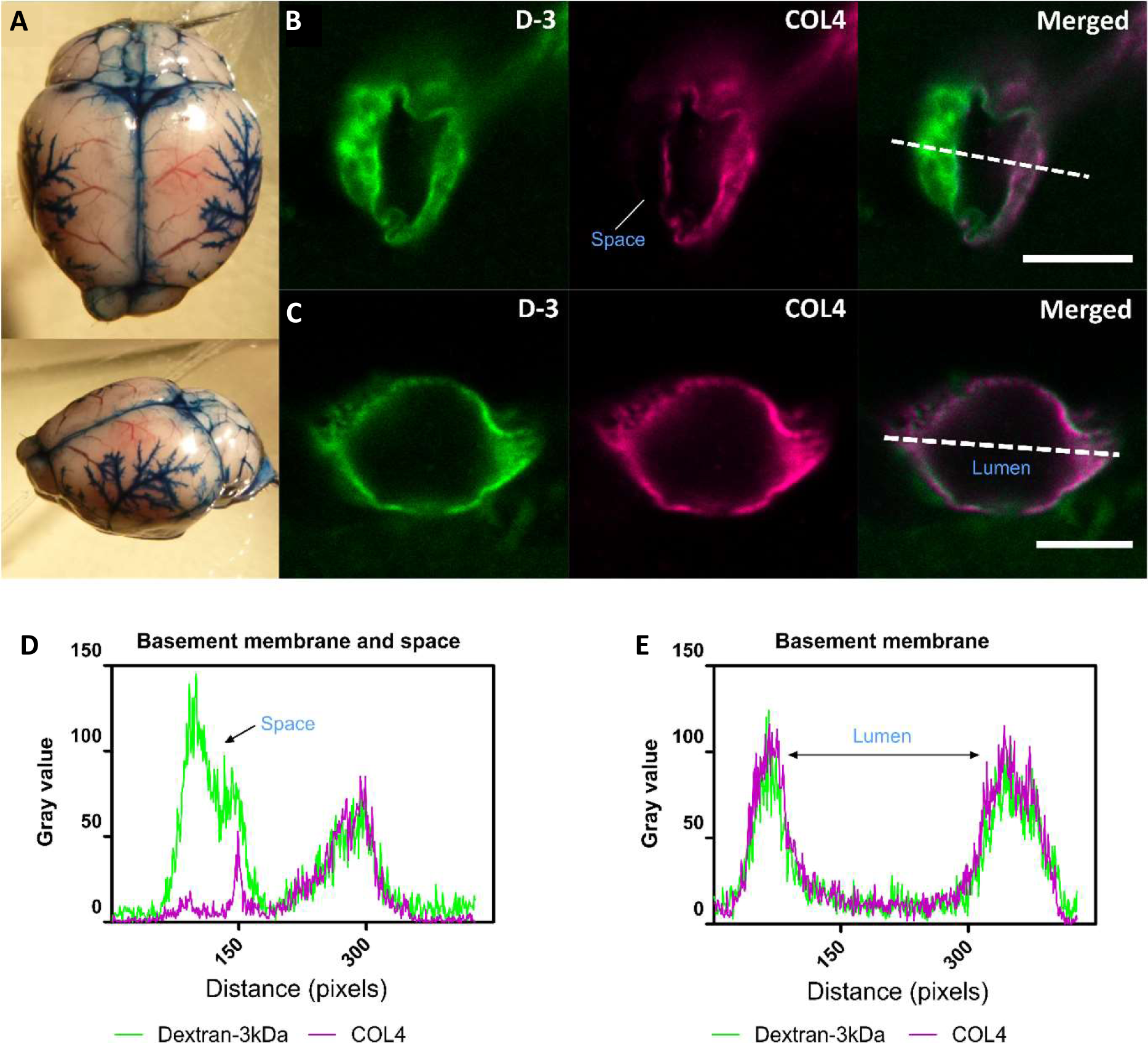
CSF tracer influx in the perivascular space. A Evans blue dye was injected into cisterna magna of a normal mouse. At the surface of the brain, dyes were found distributed along blood vessels, the middle cerebral artery (MCA) and its branches, along the superior sagittal sinus, inferior cerebral vein, and transverse sinus. B and D Co-labelling of sections with the vascular basement membrane marker COL4 revealed the localisation of CSF fluorescent tracer to the adjacent space. C and E Tracer colocalized with the basement membrane. Representative images showing spatial location of tracer soluble lysine fixable dextran 3 kDa (D-3) (green). Scale bar: top (space)=10 µm, bottom (lumen) =5 µm

### Regional CSF tracer influx is altered post-BCAS in wild-type and Tg-SwDI mice

The distribution of CSF tracer influx was then measured post-BCAS in both wild-type and Tg-SwDI mice. It was noted that the tracer distribution was quite heterogeneous between the different cohorts particularly in different brain regions, notably the dorsolateral cortex (DL CTX) and hippocampus (CA1-DG molecular layer). CSF tracer influx in the region of dorsolateral cortex was distributed along the middle cerebral artery (MCA) and its branches (Figure 3A) but this was less prominent post-BCAS (Figure 4A). Quantification of tracer indicated a significant reduction in the dorsolateral cortex post-BCAS (F (1, 27) =4.81, p=0.037) and a trend towards an effect of genotype albeit this did not reach statistical significance (p=0.064). Post-hoc analysis showed a significant reduction in WT BCAS compared to sham animals (p<0.05) (Figure 4C). It was also noted that CSF tracer was prominently distributed along the vascular network within the hippocampus but was markedly restricted post-BCAS in the hippocampal subregion: CA1-DG molecular layer (Figure 4B). It was determined that there was a reduction in tracer post-BCAS with a significant main effect of surgery (F (1, 28) =7.5, p=0.011), but no effect of genotype (p>0.05). Post-hoc tests showed a significant reduction between WT sham and BCAS mice (p=0.005) (Figure 4D). Collectively, the results demonstrate that carotid stenosis has a major impact on cortical and hippocampal glymphatic function.

**Figure 4:**
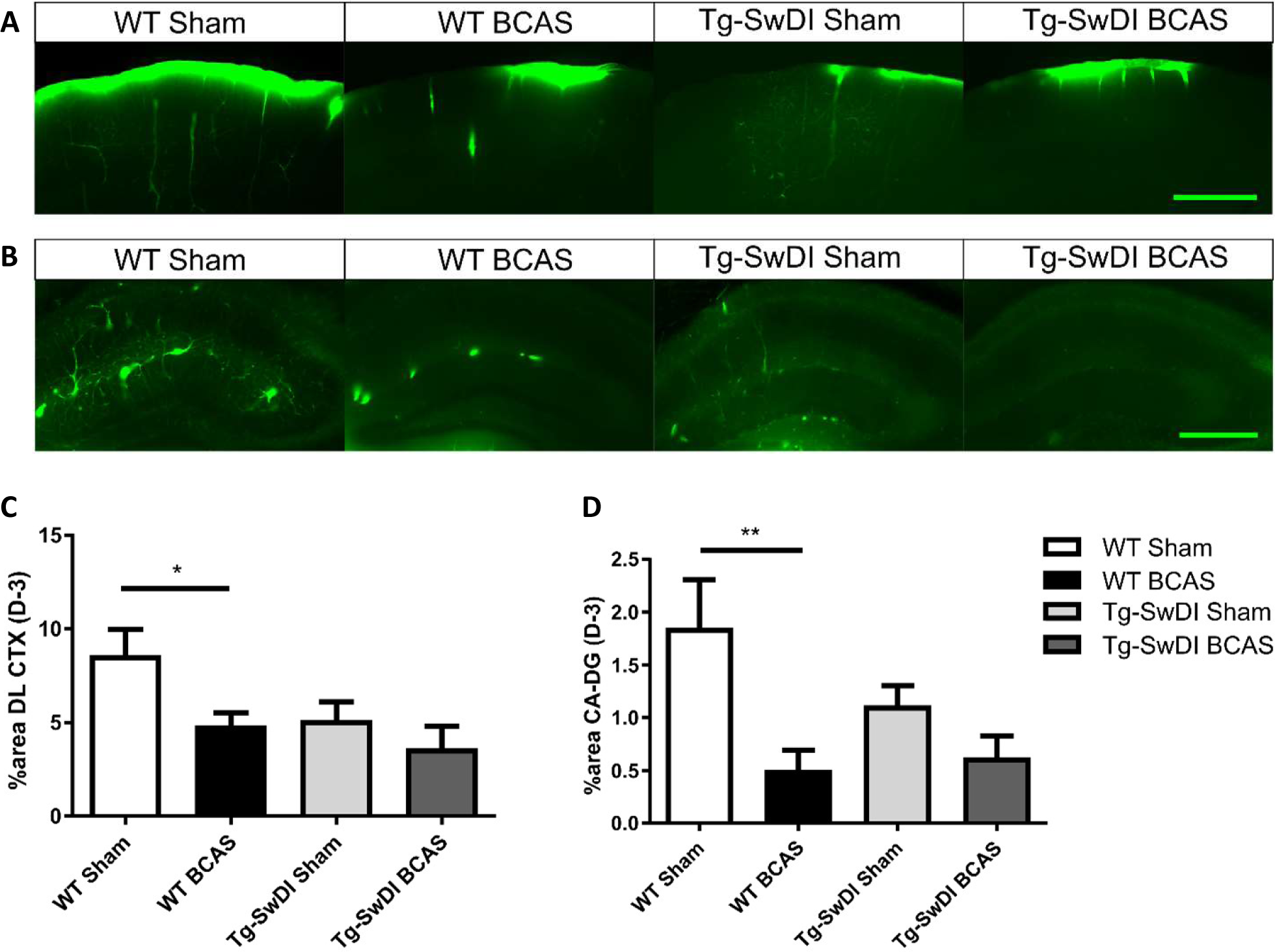
Regional CSF tracer influx is altered in BCAS and Tg-SwDI mice. Representative images of fluorescent tracer influx (D-3) (green) in the A DL CTX and B hippocampus (CA1-DG molecular layer) of WT and Tg-SwDI mice sham and post-BCAS. C and D Quantification of D-3 tracer distribution in the DL CTX and CA1-DG molecular layer. Data are presented as mean ± SEM, n=6-10 per group. Scale bar=500 µm

### BCAS exacerbates vascular amyloid accumulation

The vascular basement membranes have been proposed as pathways for the movement of fluid in the brain and involved in the build-up of amyloid causing CAA (Morris *et al.*, 2016). To investigate the potential changes of amyloid burden post-BCAS, we evaluated Aβ (6E10) load in the cortex and co-labelled with COL4 (a marker of basement membrane of blood vessels) to enable the assessment of microvascular amyloid in our Tg-SwDI mouse model with vascular amyloidosis (Figure 5A). A significant increase in the total amount of amyloid (p<0.05) and vascular amyloid was determined post-stenosis (p<0.05) in the cortex (∼250 µm from the pial surface) (Figure 5B and C, respectively). Since basement membranes have been shown as pathways for the clearance of Aβ we further determined COL4 levels but did not find significant changes post-BCAS (p>0.05) (Figure 5D).

**Figure 5:**
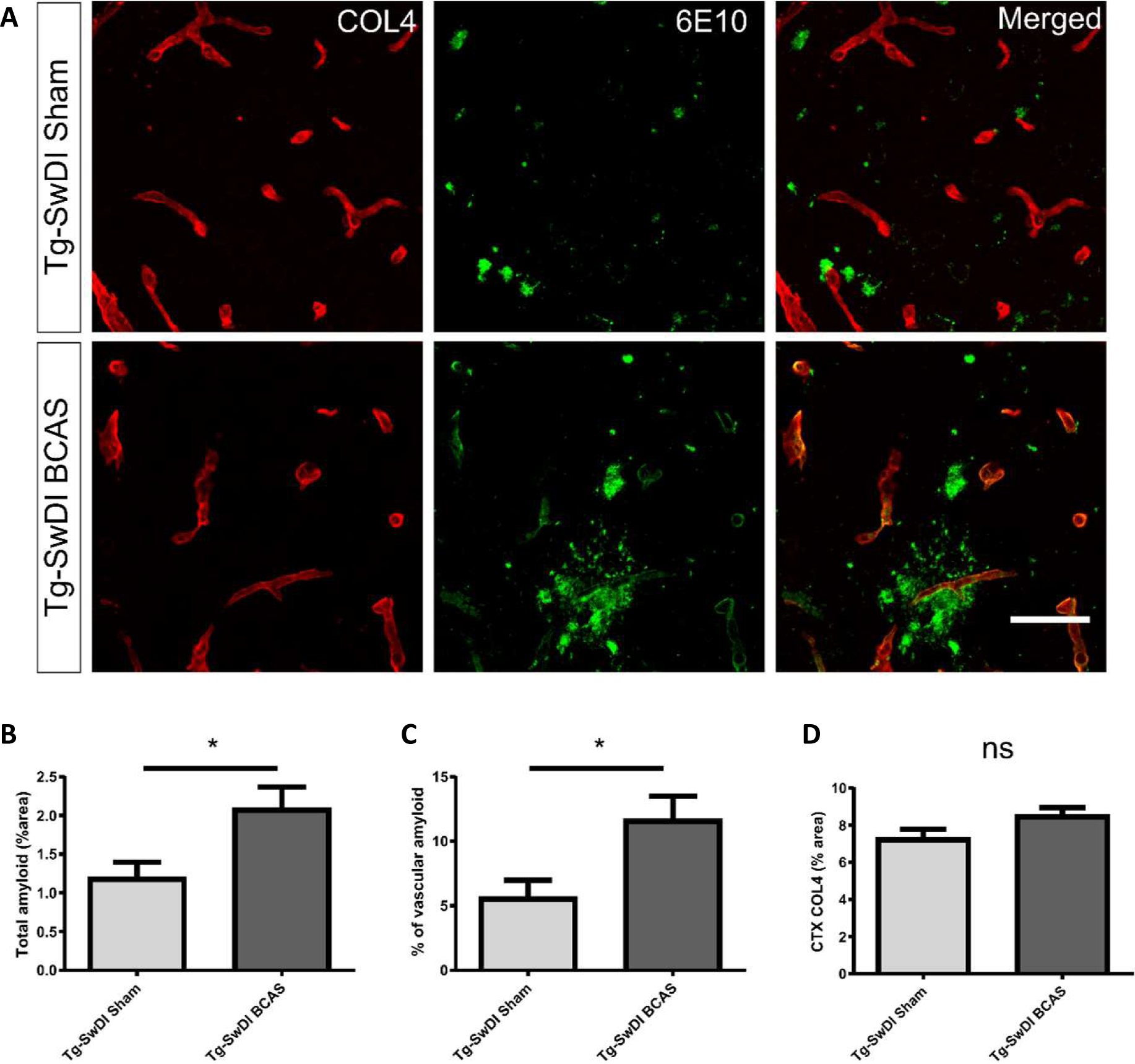
BCAS exacerbates amyloid deposition in Tg-SwDI mice. A Representative images of amyloid (green) and COL4 as a marker of vascular basement membranes (red) in the superficial brain cortex in Tg-SwDI sham and BCAS mice. B Total amyloid and C vascular amyloid were increased post-BCAS. D No significant changes of basement membrane were found. Data are mean ± SEM, n=6-9 per group. Scale bar=50 µm

### Increased astrogliosis following BCAS in cortex

To discern the mechanisms by which BCAS may impact on glymphatic function we next studied the extent of astrogliosis. Astrocytes and their end-feet have been shown to alter glymphatic function (Iliff *et al.*, 2012). GFAP immunostaining was undertaken to investigate the extent of reactive gliosis post-BCAS and in Tg-SwDI mice. BCAS surgery had a significant effect (F (1,27) =0.309, p=0.01) but there was no effect of genotype (p>0.05) on the extent of astrogliosis in the dorsolateral cortex. Post-hoc tests showed a significant increase of astrogliosis between WT sham and BCAS mice (p=0.021) (Figure 6A and C). We further analysed the hippocampal CA1- DG molecular layer. There was a significant effect of genotype (F (1, 28) =0.457, p=0.002) but no effect of BCAS (p>0.05) on astrogliosis. Post-hoc tests showed a significant increase of astrogliosis in Tg-SwDI BCAS mice compared to WT BCAS group (p=0.009) and a trend of increased astrogliosis between the WT sham and Tg-SwDI mice (p=0.061) (Figure 6B and D). Thus, alterations in astrogliosis did not always parallel the impairment in glymphatic function observed post-BCAS.

**Figure 6:**
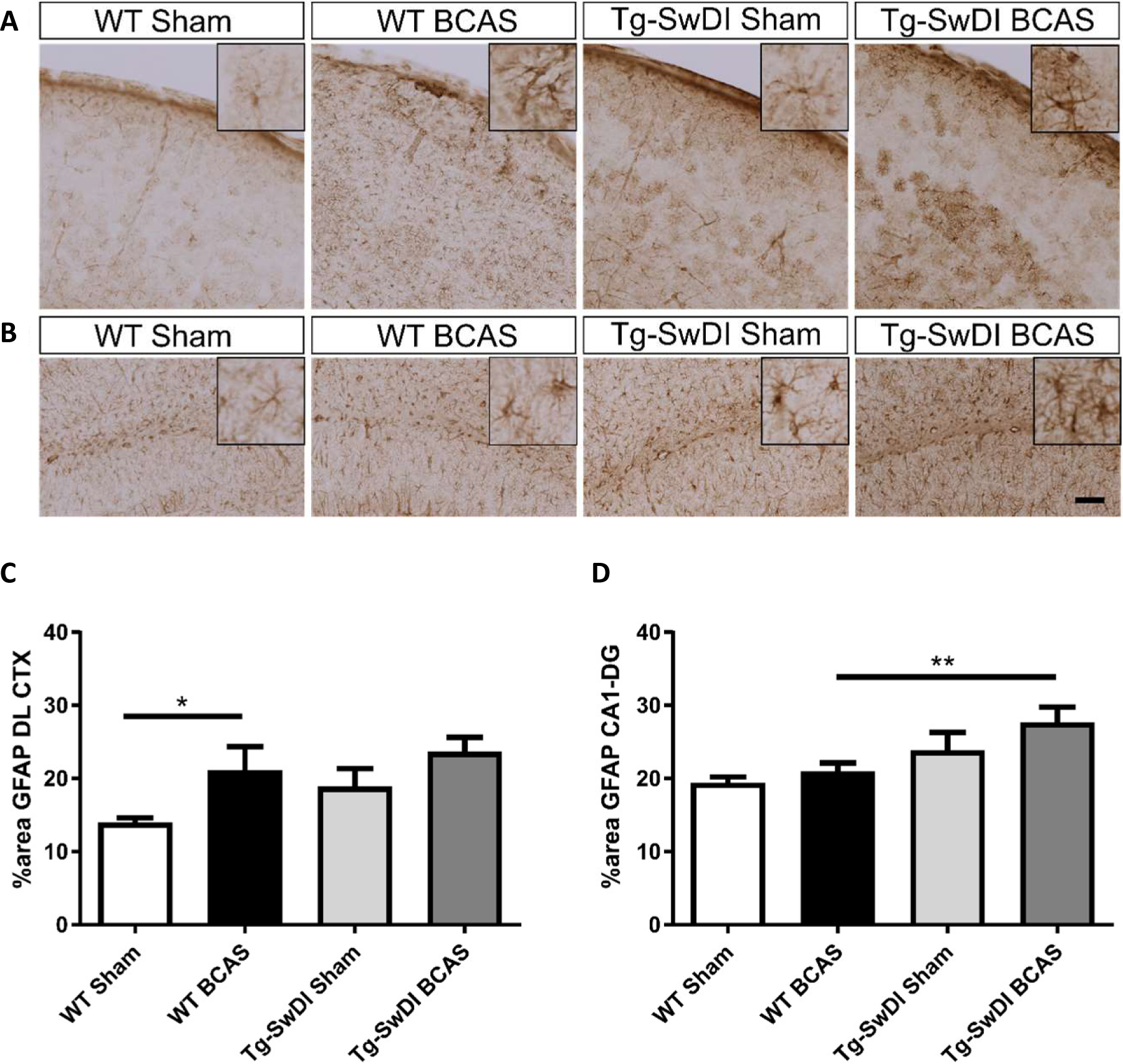
Increased astrogliosis post-BCAS in cortex. Representative images of GFAP immunostaining to assess the degree of astrogliosis in the superficial brain cortex of WT and Tg- SwDI, sham and BCAS mice. A and C In the superficial cortex, BCAS caused increased astrogliosis but was unaffected in Tg-SwDI mice. B and D In the hippocampus, there was increased astrogliosis in Tg-SwDI mice but not post-BCAS. Data are presented as mean ± SEM, n=6-10 per group. Scale bar=100 µm

### Cerebral arterial pulsation is impaired in Tg-SwDI and post-BCAS in subset of vessels

Cerebral arterial pulsation is thought to drive the glymphatic influx into and through the brain and is essential for the clearance of Aβ (Iliff *et al.*, 2013; Kress *et al.*, 2014;). Most recent evidence using two-photon imaging has supported the role of arterial pulsations in CSF movement in the perivascular spaces and basement membranes (Mestre *et al.*, 2018). To investigate whether vascular pulsation is affected in Tg-SwDI animals and post-BCAS, we used in vivo two-photon microscopy which provides high temporal resolution of individual blood vessels. We measured multiple levels of the cerebral vascular network including pial veins and arteries, penetrating arteries and ascending veins (Figure 7A). Vascular pulsatility of vessel diameter (Figure 7B) and the frequency of vessel pulsation (Figure 7C) were measured.

**Figure 7:**
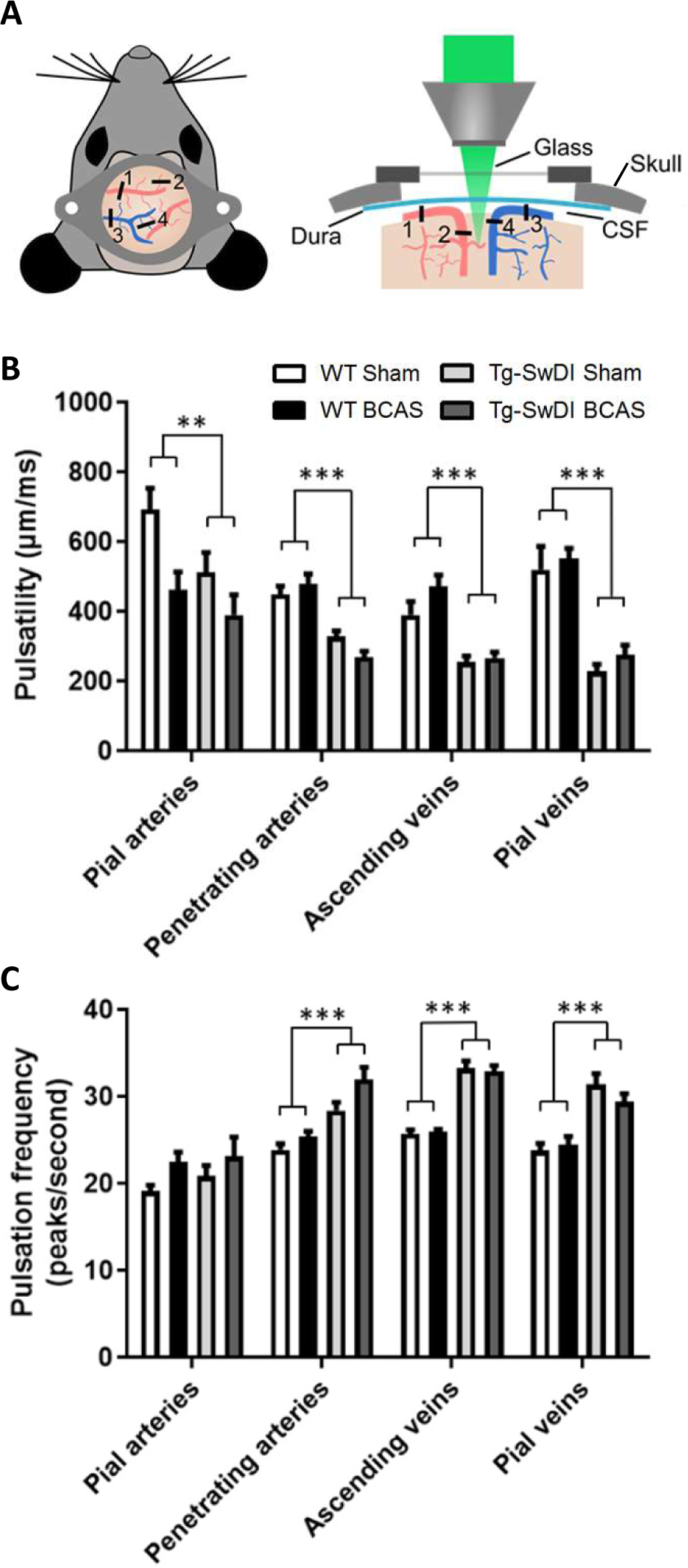
Vascular pulsatility is reduced in Tg-SwDI animals. A Two-photon microscopy was used to assess vessel pulsation in sham and post-BCAS WT and Tg-SwDI mice. Four categories of blood vessels were investigated: 1. Pial arteries; 2. Penetrating arteries; 3. Ascending veins; 4. Pial veins. B Significantly decreased pulsatility was found in pial veins, penetrating arteries and ascending veins of Tg-SwDI animals. Vascular pulsatility of pial arteries was decreased in both transgenic and BCAS animals. C There was a significant increase of frequency in the corresponding pial veins, penetrating arteries and ascending veins, but not pial arteries. BCAS caused significant increase in pulsation frequency of penetrating arteries. Data are presented as mean ± SEM, n=6-7 per group.

We found a strong effect of genotype on vascular pulsatility of vessel diameter in all types of vessels measured: transgenic animals showed reductions in pial veins (p=0.000), pial arteries (p=0.04), penetrating arteries (p=0.000) and ascending veins (p=0.000). Post hoc test revealed significant decrease in vascular pulsation between wild type sham and Tg-SwDI animals pial veins (p=0.000), pial arteries (p=0.041), penetrating arteries (p=0.001) as well as ascending veins (p=0.002). Vascular pulsation was also reduced in Tg-SwDI BCAS when compared to their wild type counterparts in pial veins (p=0.000), penetrating arteries (p=0.000), ascending veins (p=0.000), but not in pial arteries (p>0.05). Interestingly, we found significant effects of surgery in pial arteries only (p=0.005), but not in any other type of vessel. Post hoc analysis showed significant decrease in vascular pulsation in wild type BCAS animals when compared to shams (p=0.008), but no effect in Tg-SwDI animals (Figure 7B).

In addition, by measuring the frequency of pulsation, we found a significant effect of genotype in pial veins (p=0.000), penetrating arteries (p=0.000) and ascending veins (p=0.000), where the presence of transgene caused an increase in pulsation frequency. Pial arteries, on the other hand, showed no significant differences between wild-type and transgenic animals. Post hoc analysis revealed significant increase in pulsation frequency in Tg-SwDI sham animals when compared to their wild type counterparts in pial veins (p=0.000), penetrating arteries (p=0.005) and in ascending veins (p=0.000). Similarly, the pulsation frequency was also increased in Tg-SwDI BCAS animals when compared to wild type animals in pial veins (p=0.02), penetrating arteries (p=0.000) as well as ascending veins (p=0.000). BCAS surgery caused a significant increase in pulsation frequency of penetrating arteries (p=0.017), but not in any other type of vessel. Post hoc analysis here showed significant difference between Tg-SwDI BCAS and sham mice (p=0.018), but no difference in wild type animals. (Figure 7C).

## Discussion

Our findings provide experimental evidence that long-term BCAS, whilst reducing cerebral perfusion, may also affect glymphatic function, leading to cognitive impairment. This new data adds credence to a growing body of human studies that have challenged the view that reduced blood flow post-stenosis is the major contributor to VCI. Instead, alternative or additional mechanisms should be considered (Alhusaini *et al.*, 2018; Aribisala *et al.*, 2014; Shi *et al.*, 2018; Wardlaw *et al.*, 2017).

A substantial number of studies have shown that BCAS, using microcoils applied to both common carotid arteries in mice, leads to cognitive impairment (Martin *et al.*, 2016; Patel *et al.*, 2017). Consistent with these studies, we show that at 3 months after BCAS, impaired spatial learning and long-term memory are evident. The mechanistic link between BCAS and VCI has largely been attributed to the post-BCAS cerebral perfusion deficits initiating hypoxia–induced white matter pathology and degenerative changes to the glial-vascular unit (Holland *et al.*, 2015).

Our novel data suggest that BCAS can also lead to impaired CSF influx along the glymphatic pathway. In the first instance, we injected fluorescent tracer into the cisterna magna and were able to observe the tracers surrounding cerebral arteries e.g. middle cerebral artery (MCA) within the perivascular compartment (Figure 3) consistent with previous observations showing that CSF influx moves along the periarterial components into deeper brain regions (Iliff *et al.*, 2012). It has been shown by in vivo two-photon imaging that intracisternal CSF tracer travels along the perivascular component surrounding pial surface (Iliff *et al.*, 2012; Xie *et al.*, 2013). Using confocal microscopy, we were able to measure the regional distribution of tracer post-BCAS. Following 3 months of carotid stenosis impaired glymphatic function was determined in cortical and deep hippocampal regions suggesting that prolonged disruption to the vascular system may lead to enduring suppression of CSF influx to the brain.

CSF influx along the glymphatic drainage pathway has also shown to be impaired in other models relevant to cerebral vascular disease. In a rodent model of multiple infarcts, caused by intra-arterial injection of cholesterol crystals via the internal carotid artery, a transient suppression of CSF influx was determined (Wang *et al.*, 2017). However, in this study glymphatic function was restored within 2 weeks. In other models a sustained or progressive impairment of glymphatic function has been shown such as with ageing and in models relevant to AD. For example, in APP/PS1 mice, a model relevant to AD, both glymphatic periarterial influx and Aβ clearance have been found impaired, with glymphatic failure occurring prior to significant Aβ accumulation (Peng *et al.*, 2016).

To examine potential mechanisms underlying the impaired CSF influx, we assessed the extent of astrogliosis as astrocytes have an important function in CSF influx and clearance (Iliff *et al.*, 2012), and deletion of the astrocytic end feet reduces the CSF influx into the parenchyma following ischemia (Mestre *et al.* 2020). In this study there was a tendency for astrogliosis post-BCAS. In the cortex there was pronounced astrogliosis most notable in WT mice post-BCAS and in the hippocampus astrogliosis was increased in Tg-SwDI post-BCAS mice. We further explored additional mechanisms that may account for the impaired CSF influx post-BCAS. Cerebrovascular pulsatility is a key driving force facilitating CSF flow into and through brain parenchyma (Iliff *et al.*, 2013; Mestre *et al.*, 2018) and vasoconstriction was shown previously to play an important role in CSF flow following ischemia (Mestre *et al.* 2020). Using intravital imaging we found that arterial pulsation in pial vessels was affected in transgenic animals and was further exacerbated by BCAS surgery. Our results are in accordance with previous data showing that 30 minutes of unilateral ligation of internal carotid artery leads to significantly reduced pulsatility in the penetrating arteries with impaired glymphatic influx (Iliff *et al.*, 2013). Our data shows that venous pulsation, on the other hand, as well as pulsation of penetrating and ascending vessels was impaired in transgenic animals, but not affected by BCAS surgery. Together, these data suggest that BCAS has a differential effect on vascular pulsation within the vascular bed with arterial pulsation predominantly exacerbated by BCAS. Since cerebral pulsation may govern CSF-ISF exchange in murine brain leading to accumulation of solutes/proteins in the brain, this impairment may explain the accumulation of Aβ that we determined in Tg-SwDI animals post-BCAS. Altered carotid function has been associated with impaired cognitive function, greater Aβ deposition and several features of vascular disease in both human and animal studies (Ding *et al.*, 2015; Holland *et al.*, 2015; Huang *et al.*, 2012; Hughes *et al.*, 2018; Poels *et al.*, 2012). It has been also shown that the impaired glymphatic function can result in increased Aβ burden in the brain (Kress *et al.*, 2014; Peng *et al.*, 2016; Shokri-Kojori *et al.*, 2018). Another prospective population-based study has also shown alterations of pulsation in the carotid artery may contribute to the pathophysiology of cerebral microbleeds in deep brain region secondary to hypertension (Ding *et al.*, 2015). Interestingly we have also shown that sustained carotid stenosis leads to the development of vascular lesions (both microbleeds and microinfarcts) in deep subcortical structures several months post-stenosis (Holland *et al.* 2011).

The present study was restricted to evaluation of CSF influx at 3 months post-BCAS and future longitudinal approaches to evaluate potential progression of changes could be interrogated using contrast-enhanced MRI (Iliff *et al.*, 2013). In particular it will be important to discern whether glymphatic failure can be reversed and impact on the onset and progression of VCI.

## Acknowledgements

We thank R. Lennen and M. Jansen for providing technical support of MR imaging. Schematic diagrams in Figure 2 are created with Biorender.com

Author contributions (not specified as part of the submission): M. L carried out, designed and analysed most of the experiments and wrote the manuscript. A. K carried out and designed the CSF tracer experiment. J. B carried out multiphoton experiment, data analysis and some behavioural work. J. K carried out multiphoton experiment and CSF tracer experiment analysis.

J. D carried out MR imaging, behavioural and IHC experiment. B. P assisted with the behavioural study design, U. K.W assisted with the multiphoton work, R. O. C assisted the CSF tracer experiment design, data interpretation and editing manuscript. R. N. K assisted the data interpretation and editing manuscript, J. J. I assisted the study design, data interpretation and editing manuscript, K.H conducted the surgeries, supervised the project and assisted in study design, data interpretation, and writing of the manuscript. All authors assisted with editing of the manuscript.

## Funding

We gratefully acknowledge the grant support from the Alzheimer’s Society (152 (PG-157); 290 (AS-PG-15b-018); 228 (AS-DTC-2014-017)), 314 (AS –PhD-16-006), and Alzheimer’s Research

UK (ART-PG2010-3; ARUK-PG2013-22; ARUK-PG2016B-6), and The University of Edinburgh Centre for Cognitive Ageing and Cognitive Epidemiology, part of the cross council Lifelong Health and Wellbeing Initiative (G0700704/84698). M. Li and J. Beverley are funded by an

Alzheimer’s Society Scotland doctoral training programme & RS Macdonald Trust. M. Li is also funded by a China Scholarship Council (CSC)/University of Edinburgh scholarship.

## Competing interests

The authors declare that they have no competing financial interests

### Abbreviations

Amyloid-β: (Aβ)
Arterial spin labelling: (ASL)
Artificial cerebrospinal fluid: (ACSF)
Bilateral common carotid stenosis: (BCAS)
Cerebral amyloid angiopathy: (CAA)
Cerebral blood flow: (CBF)
Cerebral vascular disease: (CVD)
Dentate gyrus: (DG)
Dorsolateral cortex: (DL CTX)
Middle cerebral artery: (MCA)
Perivascular space: (PVS)
Phosphate-buffered saline: (PBS)
Region of interest: (ROI)
Subarachnoid haemorrhage: (SAH)
Transgenic mouse containing the Swedish, Dutch and Iowa mutations: (Tg-SwDI)
Vascular cognitive impairment: (VCI)
Wild-type: (WT)

Supplementary material

---

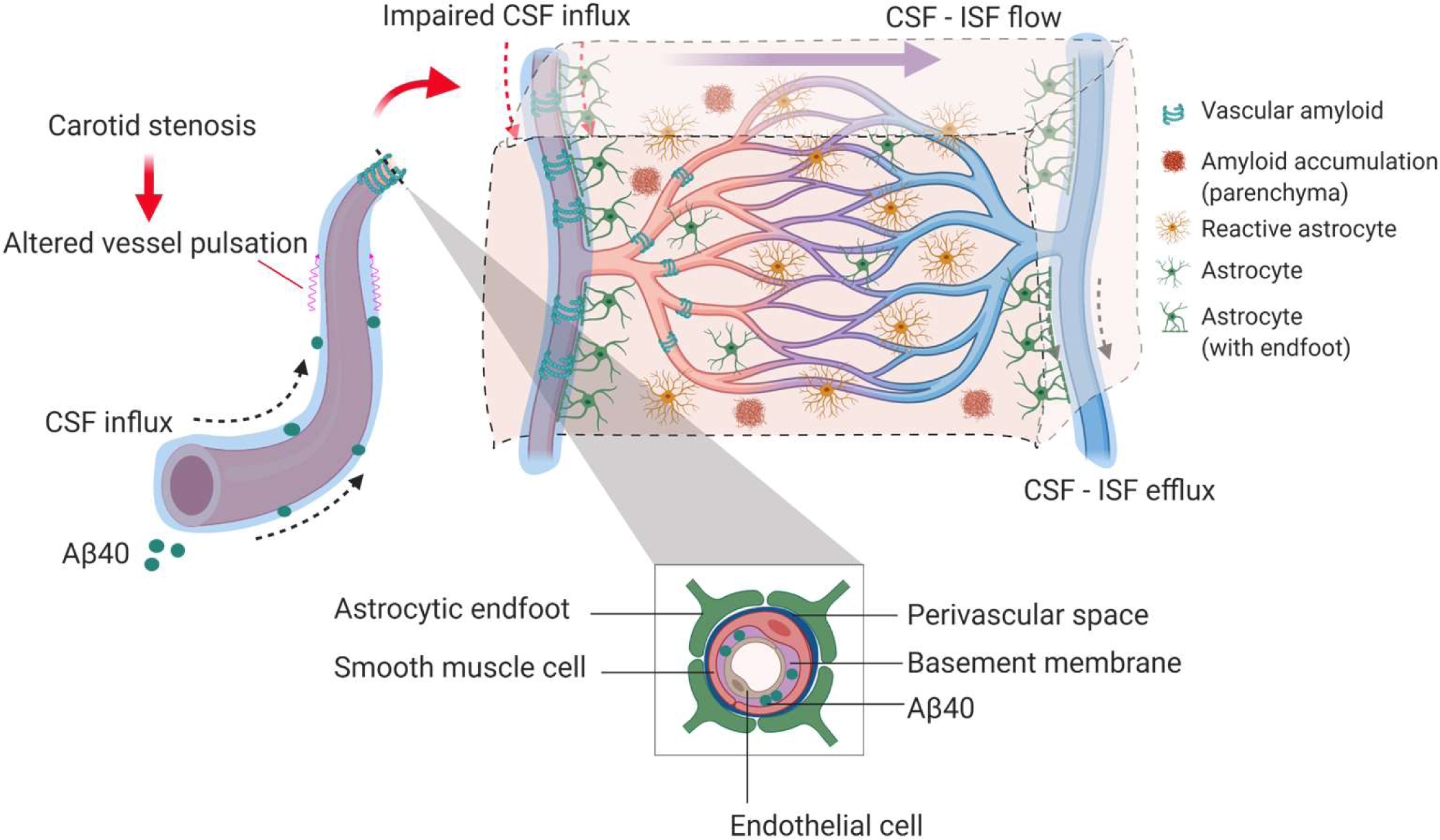

## Notes

### Competing Interest Statement

The authors have declared no competing interest.

