## Supplemental Material for "Pulsation changes link to impaired glymphatic function in a mouse model of vascular cognitive impairment"

**Title:**

Fig. S1 **Motor ability assessment in acquisition training.**

Fig. S2 **Barnes maze performance during reversal phase.**

Fig. S3 **CSF tracer influx investigated by AQP4 staining**.

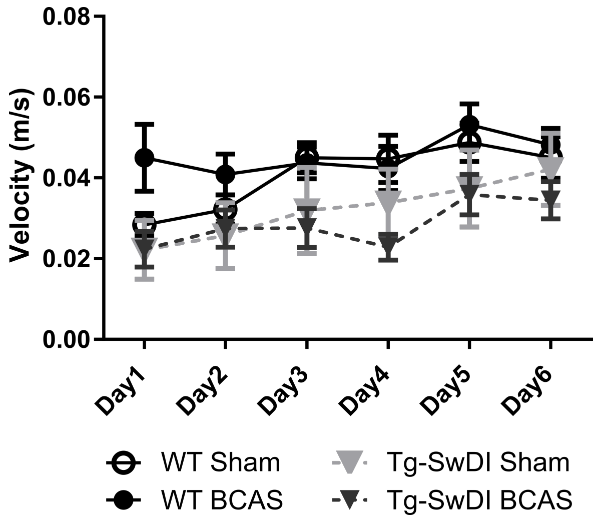


**Fig. S 1** Motor ability assessment in acquisition training. The speed or velocity was measured at the outset to assess whether motor function was affected by the genotype or surgery across different groups. There was a significant effect of genotype between WT and Tg-SwDI mice ($F(1, 30) = 8.239$, $p<0.01$) Data are mean ± SEM, n=6-10 per group

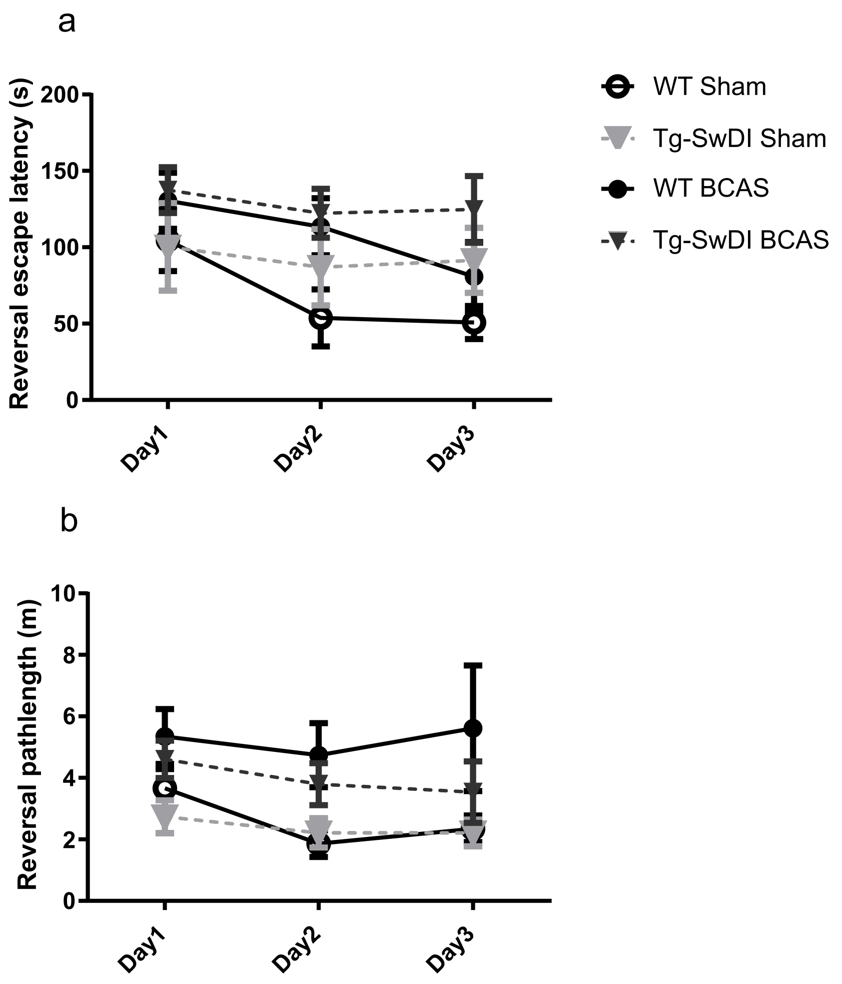


**Fig. S 2** Barnes maze performance during reversal phase. To enhance the detection of spatial learning ability, reversal trials were taken to evaluate the ability of mice to learn a new location using Barnes maze. In the reversal tests, spatial learning was assessed by comparing escape latency and pathlength over 3 days with 2 sessions per day training across all groups. a) There was a significant effect of BCAS surgery ($F(1, 30) = 4.70$, $p = 0.038$) but not genotype by comparing escape latency ($p > 0.05$). b) By comparing the pathlength in the test, there was a significant effect of BCAS surgery ($F(1, 30) = 5.80$, $p = 0.022$) but not genotype ($p > 0.05$). Data are mean ± SEM, n=6-10 per group

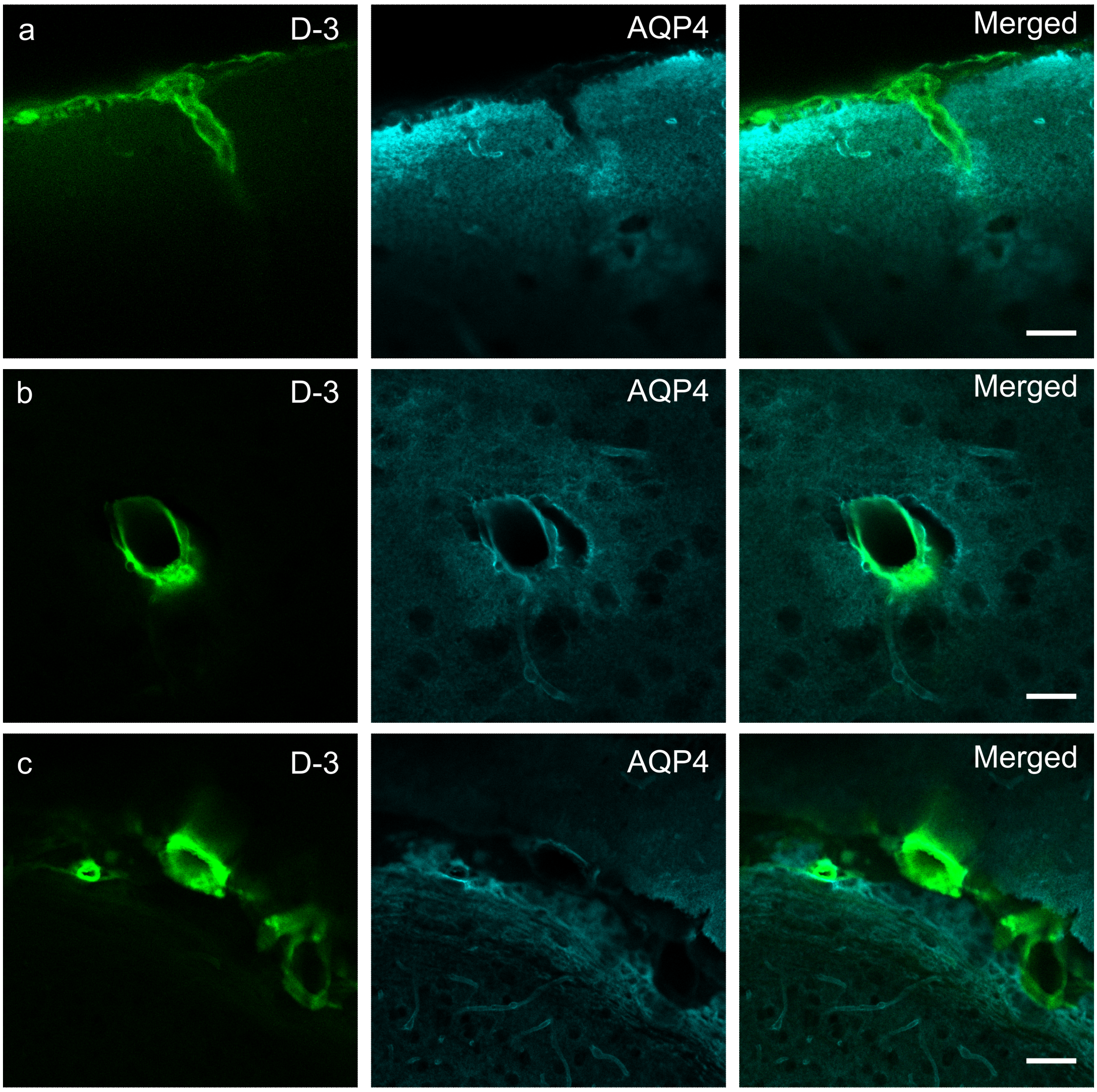


**Fig. S 3** CSF tracer influx investigated by AQP4 staining. CSF tracer Dextran 3 kDa (D-3) was surrounded by the glial astrocytic endfeet expressing AQP4. **a** Surface artery penetrating into brain surrounded by AQP4 stained brain tissue, the small tracer shows clearly filling the gap (Virchow-Robin space) at surface of brain. **b** Coronal image of superficial blood vessel and spatial location with astrocytic endfeet. **c** Tracer and AQP4 co-labelling in the deep brain region at the area between hippocampus CA3 and lateral dorsal nucleus of thalamus. Scale bars in **a-c** are 50 μm
